# Predicting spatial and seasonal patterns of wildlife-vehicle collisions in high-risk areas

**DOI:** 10.1101/2021.01.17.427044

**Authors:** Hanh K.D. Nguyen, Jessie C. Buettel, Matthew W. Fielding, Barry W. Brook

## Abstract

**Context:** Vehicle collisions with wildlife can injure or kill animals, threaten human safety, and threaten the viability of rare species. This has led to a focus in road-ecology research on identifying the key predictors of ‘road-kill’ risk, with the goal of guiding management to mitigate its impact. However, because of the complex and context-dependent nature of the causes of risk exposure, modelling road-kill data in ways that yield consistent recommendations has proven challenging.

**Aim:** Here we used a novel multi-model machine-learning approach to identify the spatio-temporal predictors, such as traffic volume, road shape, surrounding vegetation and distance to human settlements, associated with road-kill risk.

**Methods:** We collected data on the location, identity and size of each road mortality across four seasons along eight roads in southern Tasmania – a ‘road-kill hotspot’ of management concern. We focused on three large-bodied and frequently impacted crepuscular Australian marsupial herbivore species, the rufous-bellied pademelon (*Thylogale billardierii*), Bennett’s wallaby (*Macropus rufogriseus*) and the bare-nosed wombat (*Vombatus ursinus*). We fit the point-location data using ‘lasso-regularization’ of a logistic generalized linear model (LL-GLM) and out-of-bag optimization of a decision-tree-based ‘random forests’ (RF) algorithm.

**Results:** The RF model, with high-level feature interactions, yielded superior results to the linear additive model, with a RF classification accuracy of 84.8% for the 871 road-kill observations and a true skill statistic of 0.708, compared to 61.2% and 0.205 for the LL-GLM.

**Conclusions:** Forested areas with no roadside barrier fence along curved sections of road posed the highest risk to animals. Seasonally, the frequency of wildlife-vehicle collisions increased notably for females during oestrus, when they were more dispersive and so had a higher encounter rate with roads.

**Implications:** These findings illustrate the value of using data-driven approaches to predictive modelling, as well as offering a guide to practical management interventions that can mitigate road-related hazards.

## Introduction

Roads and traffic can have severe detrimental impacts on wildlife. The construction of roads can cause habitat fragmentation (Forman and Alexander 1998) and inhibit animal movement across landscapes (Trombulak and Frissell 2000). Among these impacts, wildlife-vehicle collisions could be considered the most observable, due to the visibility of the victims (roadkill). These have been known to increase mortality and therefore threaten population viability (Gibbs and Shriver 2002), with many taxa (mammals, bird, reptiles and even insects) known to be at risk of road-related injury or death (e.g., Elzanowski *et al.* 2009; Loss *et al.* 2014; Skorka 2016). Animal collisions can also cause costly damage to vehicles, and in the most extreme cases, human injury or death (Barthelmess 2014). Additionally, many generalist scavengers may benefit from roadside carrion, leading to shifts in the abundance of these species and flow-on effects within the ecosystem (Fielding *et al.* 2020; Forman and Alexander 1998). Given that the global road network will continue to expand this century there is an urgent need to understand the threat posed by roads on wildlife populations (Laurance *et al.* 2014). Quantifying and characterising the vulnerability of species and demographic components to collisions is vital for making successful management interventions (Lunney *et al.* 2008; Valero *et al.* 2015), based on both habitat and road features and all other relevant spatial and temporal components (Ramp *et al.* 2005).

In this context, we sought to demonstrate an integrative framework of: (i) road survey for wildlife-vehicle risk profiling and (ii) a complementary approach to predictive modelling, using machine learning to identify the features associated with a higher risk of wildlife-vehicle collisions using targeted field data. Linear regression has been commonly used to predict location or road-kill numbers, using physical and environmental attributes (Tejera *et al.* 2018). This method has the advantage of being simple to fit and easy to interpret (James *et al.* 2014). However, the drivers of road mortality and its spatial pattern typically include interactions and non-linearities that are not well represented with linear models (Cutler *et al.* 2007). Tree-based algorithms (e.g., random forests: Breiman 2001), which are able to handle multi-dimensional data and can selectively disregard irrelevant descriptors, are better equipped to deal with such data (Svetnik *et al.* 2003). Random forests uses out-of-bag sampling to estimate generalisation error (and thus reduce variance whilst avoiding overfitting), via random variable selection and averaged trees (Diaz-Uriarte and de Andres 2006). In this road-kill study we used random forests to deal with a mixture of categorical and continuous variables, whilst retaining an ease-of-interpretability (Bourel and Segura 2018). For comparison, we also fit the data using lasso logistic regression (Hastie *et al.* 2009), which is a generalized linear model with a coefficient shrinkage method.

Our case study was undertaken in Tasmania, the southernmost state of Australia, which is infamous for its high rate of road-kill (it is known informally as the ‘road-kill capital of Australia’: Bell 2012). Past research found that live abundance was not associated with the road-kill rate of several marsupial species within Tasmania (Nguyen *et al.* 2019). Therefore, for this study we investigated road and environmental features like traffic volume, road shape, surrounding vegetation and distance to human settlements, which are known to be associated with road-kill rates (Bond and Jones 2014; Rytwinski and Fahrig 2012). Our focal species were three iconic, large-bodied marsupials: the rufous-bellied pademelon (*Thylogale billardierii*), Bennett’s wallaby (*Macropus rufogriseus*) and the bare-nosed wombat (*Vombatus ursinus*). These animals were previously found to be the frequent victims of vehicles (based on Hobday 2010). The pademelon now is solely restricted to the island of Tasmania (it became extinct on mainland Australia in the 19^th^ century); the other two are subspecies with extant mainland counterparts. These herbivores prefer higher-quality habitats (Roger *et al.* 2007) and are still considered locally abundant (Driessen and Hocking 1992). However, they are now threatened by land-use/climate change, population suppression (e.g., culling as agricultural pests), and disease epidemics such as mange (Simpson *et al.* 2016). As such, additional road mortality poses a substantial risk to their long-term viability—and needs better contextualization with a strong evidence base for interventions.

## Methods

### Survey protocol

The road-kill monitoring was done in two rural regions of southern Tasmania that are subject to relatively high traffic volumes near the state capital of Hobart: the Huon Valley and Tasman Peninsula. Eight stretches of road (Fig. 1) were chosen for the surveys, four in each region. Based on the recommendations from previous road-kill studies in Australia (Lee *et al.* 2004; Ramp *et al.* 2006; Roger *et al.* 2011), each route was set at 15 km length (Fig. 1), covering a section of the roads with high speed limits (80–100 km/h).

**Fig. 1.**
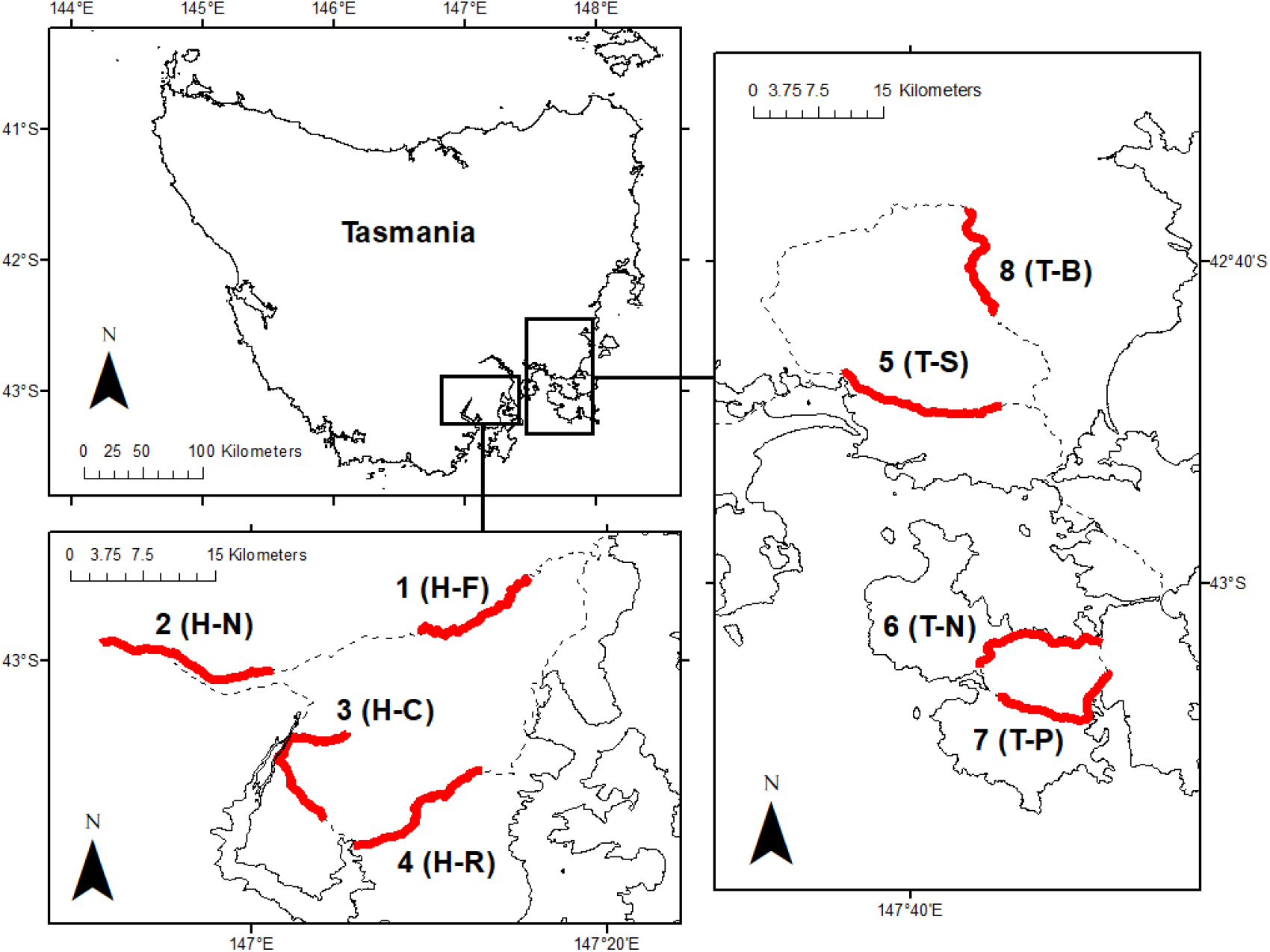
Geographic location of the road-kill surveys in southern Tasmania. The black line represents the driving route, and the red lines mark the 8 × 15 km survey roads. H = Huon Valley, T = Tasman Peninsula. 1 (H-F) = Fern Tree, 2 (H-N) = North Huon, 3 (H-C) Cygnet, 4 (H-R) = Nicholls Rivulet, 5 (T-S) = Sorell, 6 (T-N) = Nubeena, 7 (T-P) = Port Arthur, 8 (T-B) = Buckland.

Road-kill data were collected during four separate periods: September-early October 2016 (spring), January 2017 (summer), March 2017 (autumn) and July 2017 (winter). Data collection was done over a two-day period and repeated for four consecutive weeks across each season. On the first and third weeks, the routes were travelled according to their designated order (Fig. 1). The order was reversed on the second and fourth weeks, to ensure that both sides of each road were travelled equally, minimising the chance of missed observations. The roads were surveyed by two people, both spotting the animals, using a 4WD or SUV driven at low speed (~40 km/h) along the designated routes.

The vehicle was stopped for each road-kill. If it was identified as one of the target species, then the location was determined using a Garmin eTrex 30 hand-held GPS unit and recorded with a unique waypoint (WGS84 geodetic datum). Sex and species were recorded when possible. A metric tape measure was used for body, head and foot (pes) length. A separate survey was done to record road features every 50 m along the route (see *Road features as spatial predictors*). Each road-kill was labelled to permit estimation of removal rates and to avoid double-counting. Each animal observation was matched against the nearest 50 m road features at the desktop.

### Quantifying removal rates and seasonal/sex biases in frequency data

The first week of road-kill counts in each season represented bodies at various stages of decomposition, which had accumulated over an unknown prior period. Residence time on the road was dependent on both the rate of decomposition and the deliberate removal of dead animals by people or scavengers. Consequently, a high count in week one could (for instance) be attributed to many road fatalities, a low removal rate, or both. By contrast, for weeks two to four, both the number of new additions and the disappearance of previously recorded bodies during the past week were known (because each specimen was individually tagged/geo-located), resulting in direct information on accumulation and removal rates for the periods spanning weeks 1–2, 2–3 and 3–4, for each season. These data enabled us to estimate the number of animals that were never observed (i.e., those killed but subsequently removed *between* sampling times). Taken together, these multiple lines of evidence on biases allowed for the calculation of a re-calibrated count for each season (i.e., adjusted for weekly variation in removal rates and missing observations). Contingency tables were used to summarise sex biases in road-kill across four seasons for the three species. Differences in category counts were evaluated statistically using Fisher’s exact binomial test (Suissa and Shuster 1985), with an expected (null) sex ratio of 50 %. A chi-squared test was used to determine whether the number of road-kill were distributed randomly across seasons (Rorden *et al.* 2007).

### Data specification for spatial analysis

The dependent variable for the modelling was presence or absence of wildlife fatalities at a given location, with the predictors being both categorical (habitat, road and fence) and continuous (human population density (pop) and distance to fresh water (DFW). Habitat was classified as either FF = forest on both sides of the road, FO = forest on one side, open on the other, or OO = open on both sides. Road shape was designated ‘straight’ = S or ‘curved’ = C, and a roadside verge fence was noted as present or absent. Regularly spaced background points were generated at 50 m intervals along the designated routes, using ArcGIS on a shapefile of Tasmanian roads, which resulted in a total of 2, 491 points. Each regular point was assessed (both on-ground and via examination of high-resolution satellite imagery via Google Earth) for habitat and road characteristics. Human population density was extracted from data obtained from the Center for International Earth Science Information Network (ciesin.org) and distance to water was processed in ArcGIS 10.4. To use as (pseudo)absence points in the modelling, the regular points were then filtered by (conservatively) only including the subset of points spaced at least 200 m from any given road-kill record. This approach yielded a total data set of 871 road-kill-presence points and 674 pseudoabsences.

### Statistical analysis of spatial predictors of wildlife-vehicle collisions

Machine-learning algorithms were run using (i) random forests fit using R 3.5 library party (Hothorn *et al.* 2006) and (ii) lasso-logistic regression fit using glmnet (Friedman *et al.* 2010). Random forests (Strobl *et al.* 2007) can be used as a binary classifier with interactions: branch splitting is based on the average of many random subsets of predictors, with the goal of decorrelating the individual trees. However, like other tree-based machine-learning methods, the random forests algorithm can be prone to bias in variable selection if categorical variables with many levels are included (Torsten *et al.* 2006). To mitigate this issue and properly account for a model structure that considers both continuous and categorical predictors that are uncorrelated, the use of an unbiased tree algorithm is recommended, along with permutation variable importance (Augustin *et al.* 2008; Strobl *et al.* 2007). For easy interpretation, we also used partial dependence plots to illustrate the relationship between the predictors and the roadkill (Visintin *et al.* 2016).

To evaluate the predictive power of random forests against linear models, we also fitted the data using an additive logistic regression and selected the appropriate model components and dimension using machine learning, via a lasso (least absolute shrinkage and selection operator) regularization analysis (Hastie *et al.* 2009). The lasso estimates an optimally scaled penalty to be applied to model coefficients, resulting in sparse models that are not overfit. This is akin to continuous subset selection, whereby the shrinkage penalty (λ)—determined adaptively by repeated k-fold cross-validation—is chosen to minimize out-of-sample prediction error. Model adequacy was measured based on classification accuracy, sensitivity, specificity and the true skill statistic (balancing errors of omission and commission: Allouche *et al.* 2006).

## Results

### Seasonal and sex differences in road-kill risk

The most frequently road-killed species was the pademelon (Table 1). The raw counts of road-kills differed substantially across seasons: 201 in spring, 111 in summer, 256 in autumn and 301 in winter. Road-kills persisted longest on roads in spring and disappeared most rapidly in summer, leading to re-calibrated bias-corrected counts: 124 in spring, 178 in summer, 266 in autumn and 303 in winter. The calibrated results showed no significant difference in road-kill accumulation rates between winter and autumn (95% CI of exact binomial test = 0.426–0.509; overlaps 0.5 null expectation), while spring had the lowest seasonal rate (autumn : spring ratio = 0.684, CI = 0.635–0.730; summer : spring = 0.589, CI = 0.532–0.645; winter : spring = 0.710, CI = 0.664–0.752). Autumn and winter remained the seasons with the highest road-kill for macropods post-calibration.

**Table 1.** Total road-kill frequency (counts) of the rufous-bellied pademelon (*Thylogale billardierii*), Bennett’s wallaby (*Macropus rufogriseus*) and bare-nosed wombat (*Vombatus ursinus*) surveyed along 8 × 15 km roads across four weeks within each season (spring, summer, autumn and winter) of southern Tasmania, during 2016-2017. Unsexed animals were too macerated or dismembered to permit assignment of a gender.

| Species | Male | Female | Unsexed | Total |
| --- | --- | --- | --- | --- |
| *T. billardieri* | 261 | 170 | 217 | 648 |
| *M. rufogriseus* | 63 | 52 | 63 | 178 |
| *V. ursinus* | 10 | 14 | 19 | 43 |

There was no overall sex bias in road-kill for the wallaby (0.452, CI = 0.359-0.548) but a slight male bias for the pademelon (0.606, CI = 0.558-0.652). Female pademelons were most at risk in autumn (Fig. 2a), whereas winter was the only season with a male bias (0.744, CI = 0.668-0.810). For the wallaby, female mortality was highest in autumn (0.713, CI = 0.610-0.801), and this persisted to winter (Fig. 2b), with no clear peak for male-wallaby mortality, although their numbers decreased noticeably in summer and then gradually recovered (Fig. 2a, b) with a significant male bias in spring (0.867, CI = 0.693-0.962). There was no statistically meaningful sex bias or trend for the wombat due to sparse numbers (Fig. 2c).

**Fig. 2.**
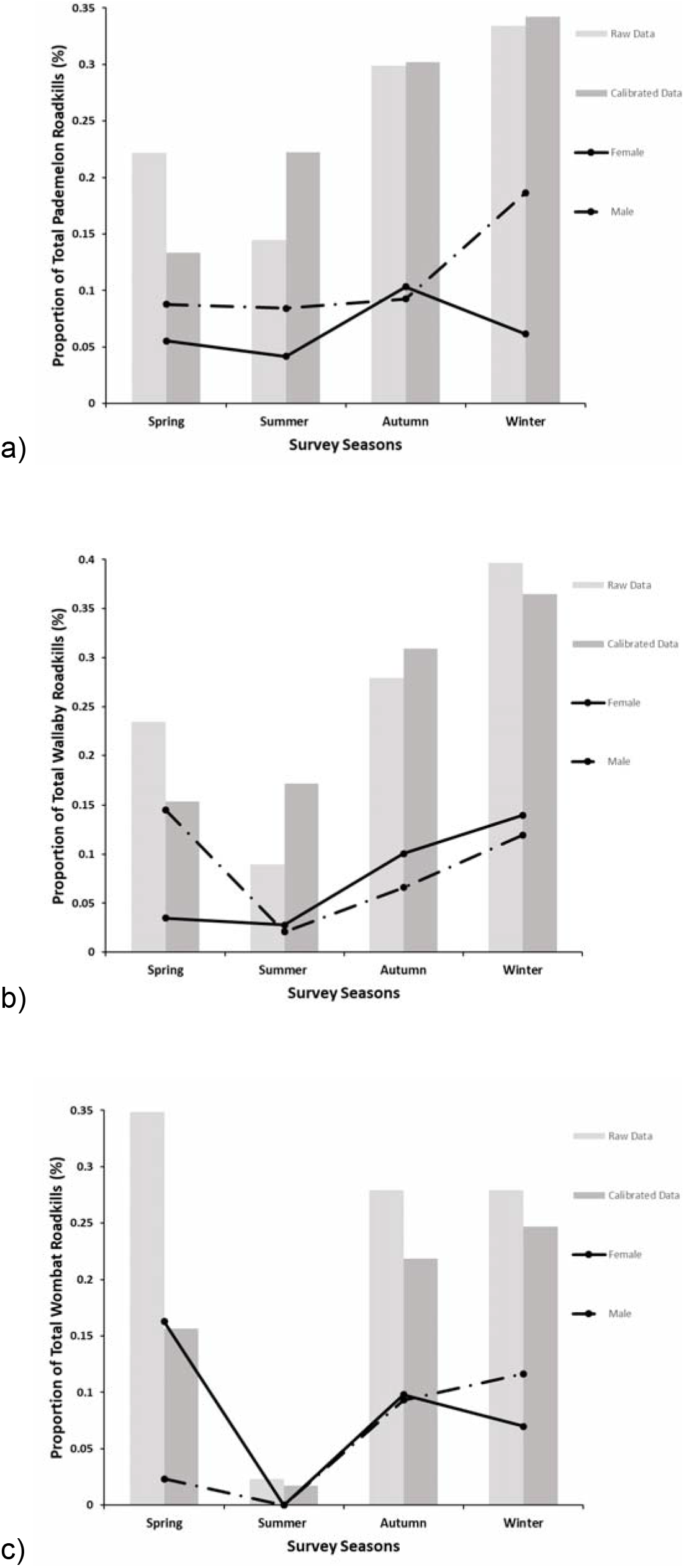
Seasonal variation in total abundance and sex bias of marsupial road-kills in southern Tasmania in 2016-2017, showing results before and after the observational calibration (to account for removals and missed observations). Plotted are: a) *Thylogale billardierii*, b) *Macropus rufogriseus* and c) *Vombatus ursinus*. Raw and calibrated bars are expressed as each season’s fraction of the total count; lines show the proportion of each sex found in a given season relative to all seasons.

### Spatial predictors of road-kill

The random forests algorithm produced a minimized out-of-bag error when three of the predictor variables were randomly sampled as candidates at each branch split. The most consistently influential variable in the decision trees was habitat, with an importance value of 0.146. The two continuous predictors (human population density and distance to fresh water) were the second-most selected across the tree-averaging process, yielding variable importance values of 0.117 and 0.114 respectively. The highest risk of road mortality was associated with forested areas, in locations with no fence and along curved sections of road (Fig. 3). The complex relationship between the continuous variables to road-kill presence is shown in Fig. 3. The overall classification accuracy of the random forests model of road-kill risk was 84.8%: 91.3% for presences (76 errors in 871 observations) and 76.4% for pseudoabsences (159/674), with a sensitivity = 0.833, specificity = 0.871, and true skill statistic (TSS) = 0.704 (where TSS = 1 is a perfect classifier, and 0 the random expectation).

**Fig 3.**
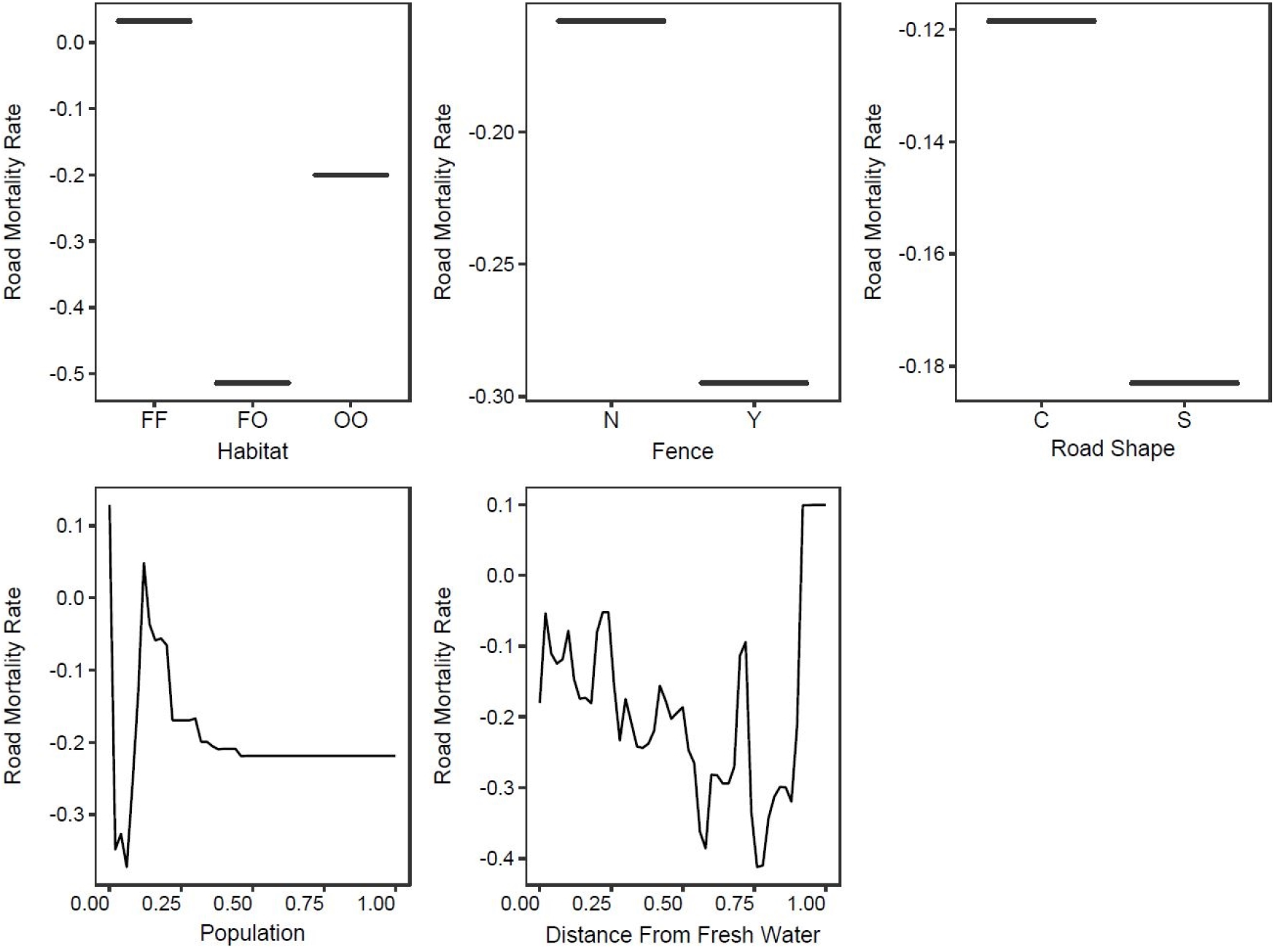
Partial dependence plots of the predictors on the road-kill rate of the three Tasmanian marsupials, derived from a machine-learning algorithm fitted using random forests.

For comparison, the lasso-based shrinkage of a logistic regression model rejected all but two categorical predictors to derive the most parsimonious model (at λ = 0.024; Fig. 4): habitat (coefficients relative to FF were: β_FO_ = 0.613 and β_OO_ = 0.063) and fence (coefficient relative to no fence, β_fence_ = 0.470). As such, risk of vehicle fatality in this linear-additive model was predicted to be highest in habitats with a mix of forest and open areas with the road bordered by fences. The lasso-model classification accuracy for road-kill location was, however, markedly lower than that for the random forests, at 61.2% overall: 76.1% for presences and 42.0% for pseudoabsences, with a sensitivity = 0.629, specificity = 0.576, and true skill statistic = 0.205 (i.e., a bias towards over-estimating the likelihood of road-kill at a location).

**Fig 4.**
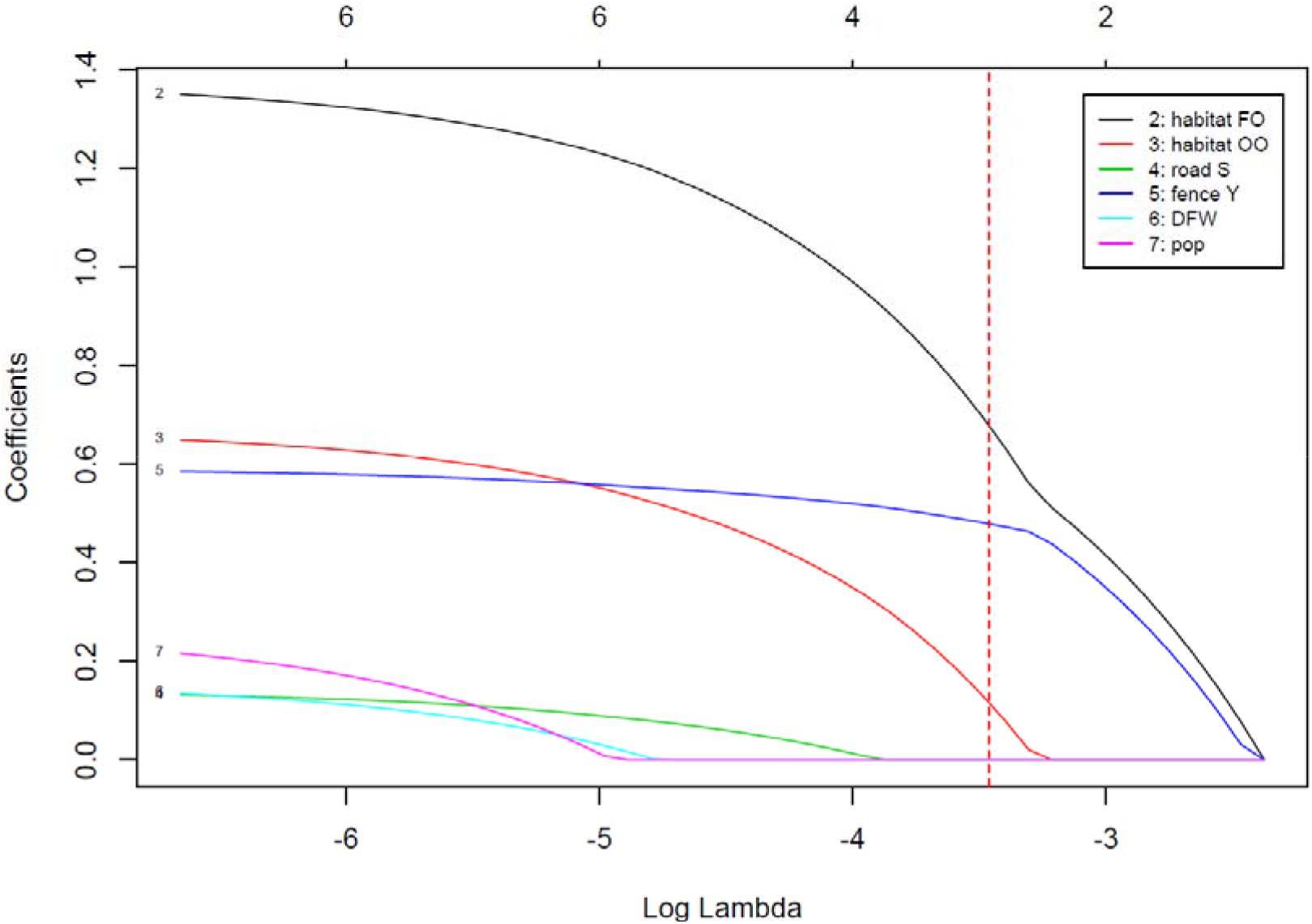
Lasso-based regularization of an additive logistic-regression model, showing changes in the model coefficients with respect to the lasso-shrinkage parameter (lambda); also shown are number of retained coefficients (above the plot window). The dashed vertical red line is the value of lambda that minimized the cross-validation error, which resulted in a sparse three-parameter model of habitat (forest-open, open-open) and fence (present) relative to the forest-forest/fence-absent base model. All other predictors were shrunk to zero.

## Discussion

### Machine-learning approaches to spatial analysis of road-kill risk

In this road-kill study, the random forests classification algorithm was clearly superior to lasso-logistic model for predicting the riskiness of locations for wildlife-vehicle collisions. The lasso-regularization of a generalized linear model disregarded both continuous variables (distance to freshwater and population), retaining only two categorical variables: habitat and fence. However, as illustrated by the partial dependence plots (Fig. 3), distance to freshwater and population clearly displayed a non-linear relationship to road mortality, which explained why they could not be captured in the additive logistic model. This model-algorithm comparison shows the value of tree-based machine-learning approaches to developing more complex (yet parsimonious and data-driven) models for analysing road mortality data with non-linearities are plausible.

Additionally, the random forests approach is able to implicitly account for feature interactions in the data without the need to manually identify and include specific terms in the model, thus reducing the need for hand-crafted *a priori* specification (Thuiller *et al.* 2009; Wright *et al.* 2016) of interactions. On the other hand, random forests is less effective in dealing with linear relationships, and linear models can be better at interpolating data (Thuiller *et al.* 2009). Furthermore, the output of random forests is comparatively harder to interpret than that of generalized linear models, becoming more complicated as the number of nodes increases. While partial dependence plots are useful for visualising results, they cannot be used to interpret interactions (Cutler *et al.* 2007). Therefore, prior to selecting the appropriate analysis, it is crucial that the purpose of the study be considered. Is it to analyse the most important predictors for road-kill? Is it to extrapolate the data and predict the rate of road-kill in similar areas? This decision can guide the appropriateness of the method(s) selected for modelling.

Regarding the specific results of this case study in Tasmania, the macropod species leave their diurnal shelters, located in closed habitats (e.g., native forests, established plantations), to forage in more open adjacent areas (e.g., grassland, farmland, young plantations) at night (le Mar and McArthur 2005; Wiggins and Bowman 2011). Wombats also prefer open high-quality grazing land adjacent to cover (Evans *et al.* 2006). Given these behaviours, half-forested areas should intuitively have the highest rates of road-kill (and this was predicted by the lasso-logistic). However, when interactions are considered (in random forests), forested areas were found be at highest risk. One explanation might be that forested areas were typically located further from populated areas, leading to changes in driver attitude as speed limits increased (based on Hobday 2010). It is likely that animal behaviour (e.g., foraging routes, preference for well-vegetated roadside verges) would also change with regards to proximity to fresh water and availability of pastures across mixed-agricultural lands.

Most of the surveyed routes adjacent to farmland and plantations were fenced off using stock races, which has been found to correlate positively with the frequency of kangaroo-vehicle collisions (Lee *et al.* 2004). The directional effect of fencing on the frequency of animal-vehicle collisions have been found to be both positive and negative, often depending on its type, suitability to the target-species ecology, and the state of maintenance (Polak *et al.* 2014). Therefore, a fencing-to-mitigate strategy might not be appropriate if the target-species have widely different ecologies, lowering risk for one but increasing it for another.

### Seasonal and autecological influences on road-kill rates

The fewest road-kills were seen in summer, for all species. This result appears to agree with the hypothesis that a longer day length reduces the number of animal-vehicle collisions, because the largely crepuscular/nocturnal habits of Australian mammals lead to most road-kills occurring during the dusk-to-dawn period, when animals have commenced foraging but vehicle activity is still relatively high (Ellis *et al.* 2016). However, we were able to estimate higher decomposition rates of summer, likely due to warmer temperatures and higher insect abundance (Campobasso *et al.* 2001), along with the more rapid removal of roadkill by roadside-management organisations during peak tourist seasons. After taking into consideration this higher rate of road-kill removal via data calibration, our results revealed no difference between spring and summer collision frequency for the wallaby, with a slightly higher rate in summer compared to spring for the pademelon. As such, the nocturnal-overlap-risk hypothesis is only weakly supported by our corrected data.

The striking increase in female counts in autumn for the macropod species coincides with the peak period of parturition (Curlewis 1989). Female macropods in oestrous tend to increase their mobility to enhance their exposure to males (Fisher and Lara 1999). Another study showed that female wallabies in southern Australia also extended their home range in mid-year (when measured road-kill was high), due to harsher weather driving them to move to access more resources (Johnson 1987). Consequently, their chance encounter rate with roads is likely to be higher over this period, leading to more vehicle collisions. By contrast, in spring and summer, females of both species tend to restrict their movements when joeys leave the pouch (Rubenstein and Wrangham 1986), resulting in lower road-kill counts. The road-kill analysis of the bare-nosed wombats (*Vombatus ursinus*) was constrained due to the small and regionally restricted sample sizes, prohibiting any robust interpretation of trends.

### Requirements for further targeted research

Past road-kill research has faced an obvious trade-off. Studies that survey multiple stretches of roads had robust spatial coverage but limited temporal implications (Mallick *et al.* 1998; Russell *et al.* 2009), whereas those that lacked an explicit spatial contrast usually compensated with a more frequent and consistent approach to surveying (Ramp *et al.* 2005; Roger *et al.* 2011). Our approach sought to reach an acceptable middle ground: repeatedly surveying enough short segments of roads to yield a broad spatial coverage across all seasons, along with marking and geolocating the wildlife corpses to permit explicit estimation of biases and improve the overall systematic rigor of the surveys. We have provided our data and the random forests predictive model to the relevant wildlife and road-management authorities in Tasmania, to help guide future planning and decision making.

The ultimate goal of investigating road mortality in animals—besides determining correlating/causal factors and using these for prediction and intervention—is to understand any possible effect that this additive death rate has on absolute population viability, particularly for species threatened by other risks (Roger *et al.* 2011). An increase in the loss of female macropods during oestrous in autumn could have fitness implications for recruitment in local populations, especially if new threats (e.g., accidental introduction of the red fox, *Vulpes vulpes*) threaten future demographic stability. However, this effect, and any similar sex biases or seasonal trends, could only be reasonably used to predict the effect of road mortality on populations when other autecological information are well understood (Ramp *et al.* 2005). For (most) situations in road ecology, detailed demographic data on the underpinnings of population viability (e.g., density-dependence, habitat suitability, age-specific vital rates), are lacking, making it difficult to determine additive or compensatory effects of vehicle-collision mortality, especially as an interaction with other drivers of population dynamics. A future synergy of mechanistic viability modelling and targeted field studies, spanning space (e.g., near and distant from roads) and time, is clearly ideal. That said, we have here demonstrated that advanced approaches to the modelling of pattern-based information on road-kill can still lead to useful and nuanced insights on relative risk, and thereby offer a guide to practical management interventions that can mitigate these hazards.

## Acknowledgements

We would like to thank Dr. Emily Flies, Dr Stefania Ondei and Dr Sanghyun Hong for their valuable inputs. Research was funded by ARC grant FL160100101. Road-kill data were collected under the University of Tasmania animal ethics permit A0015929.

